# Virological characteristics of the SARS-CoV-2 BA.2.86 variant

**DOI:** 10.1101/2023.11.02.565304

**Authors:** Tomokazu Tamura, Keita Mizuma, Hesham Nasser, Sayaka Deguchi, Miguel Padilla-Blanco, Keiya Uriu, Jarel Elgin M. Tolentino, Shuhei Tsujino, Rigel Suzuki, Isshu Kojima, Naganori Nao, Ryo Shimizu, Michael Jonathan, Yusuke Kosugi, Ziyi Guo, Alfredo A Hinay, Olivia Putri, Yoonjin Kim, Yuri L Tanaka, Hiroyuki Asakura, Mami Nagashima, Kenji Sadamasu, Kazuhisa Yoshimura, The Genotype to Phenotype Japan (G2P-Japan) Consortium, Akatsuki Saito, Jumpei Ito, Takashi Irie, Jiri Zahradnik, Terumasa Ikeda, Kazuo Takayama, Keita Matsuno, Takasuke Fukuhara, Kei Sato

**Author notes:** Corresponding authors (Jiri Zahradnik), (Terumasa Ikeda), (Kazuo Takayama), (Keita Matsuno), (Takasuke Fukuhara), (Kei Sato). These authors contributed equally (Tomokazu Tamura, Keita Mizuma, Hesham Nasser, Sayaka Deguchi, Miguel Padilla-Blanco, Keiya Uriu, Jarel Elgin M. Tolentino).

## Abstract

In late 2023, a lineage of SARS-CoV-2 emerged and was named the BA.2.86 variant. BA.2.86 is phylogenetically distinct from other Omicron sublineages identified so far, displaying an accumulation of over 30 amino acid mutations in its spike protein. Here, we performed multiscale investigations to reveal the virological characteristics of the BA.2.86 variant. Our epidemic dynamics modeling suggested that the relative reproduction number of BA.2.86 is significantly higher than that of EG.5.1. Experimental studies showed that four clinically-available antivirals were effective against BA.2.86. Although the fusogenicity of BA.2.86 spike is similar to that of the parental BA.2 spike, the intrinsic pathogenicity of BA.2.86 in hamsters was significantly lower than that of BA.2. Since the growth kinetics of BA.2.86 is significantly lower than that of BA.2 in both *in vitro* cell cultures and *in vivo*, it is suggested that the attenuated pathogenicity of BA.2.86 is due to its decreased replication capacity.

## Introduction

In November 2023, the SARS-CoV-2 XBB descendants, mainly EG.5.1 (originally XBB.1.9.2.5.1), are predominantly circulating worldwide according to Nextstrain (https://nextstrain.org/ncov/gisaid/global/6m). However, a lineage far distinct from XBB unexpectedly emerged and named BA.2.86 on August 14, 2023 ^1^. Notably, BA.2.86 bears more than 30 mutations in the spike (S) protein compared to XBB and the parental BA.2, which are assumed to be associated with immune evasion ^2^. According to the higher number of amino acid substitutions in this variant, the WHO immediately designated BA.2.86 as a variant under monitoring on 17 August 2023 ^3^. As of October 31, 2023, the BA.2.86 variant has been identified globally, with an increasing frequency in viral genome surveillance. To date, over 1,400 sequences of BA.2.86 and its related lineages have been reported on GISAID (https://gisaid.org/).

The immune evasive potential of BA.2.86 has been evaluated in recent studies including ours ^2,4–9^. Additionally, some studies addressed the virological features of BA.2.86, such as the affinity of the receptor-binding domain (RBD) of the BA.2.86 S to ACE2 receptor ^5,8^ and the fusogenicity of BA.2.86 S ^7^. Moreover, Khan et al. have recently reported the growth kinetics of a clinically isolated BA.2.86 live virus in *in vitro* cell cultures ^7^. However, the sensitivity of BA.2.86 to clinically available antiviral drugs and the intrinsic pathogenicity of BA.2.86 in hamsters remain unknown. Here, we elucidated the virological features of a newly emerging SARS-CoV-2 BA.2.86 variant.

## Results and Discussion

### Epidemic dynamics of BA.2.86

BA.2.86 is phylogenetically distinct from other Omicron sublineages that have emerged to date, exhibiting the accumulation of over 30 amino acid mutations in the S protein (**Figures 1A and S1A**). This indicates that BA.2.86 may have unique characteristics when compared to other Omicron subvariants. To assess the epidemic potential of BA.2.86, we estimated its relative effective reproduction number (R_e_) (**Figures 1B, 1C, and S1B and Table S1**). Although we previously estimated the R_e_ of BA.2.86, that study did not conclusively determine whether BA.2.86 shows significantly higher R_e_ than EG.5.1, the currently dominant lineage globally ^2^. This was mainly due to the considerable uncertainty on the estimated R_e_ of BA.2.86, attributed to a limited sequence dataset at that time. In our current analysis, we more accurately estimated the relative R_e_ of BA.2.86 by incorporating genome surveillance data from six countries where BA.2.86 is proliferating using a Bayesian hierarchical multinomial logistic model ^10,11^]. This method enabled us to estimate the R_e_ of each variant within individual countries (**Figure S1B**) as well as a global R_e_ average (**Figure 1B**). We show that the global average R_e_ of BA.2.86 is 1.07-fold higher than EG.5.1 (**Figure 1B**). Additionally, in each country examined, the R_e_ of BA.2.86 significantly exceeded that of EG.5.1 (**Figure S1B**). Indeed, BA.2.86 is gradually growing in European countries such as Denmark, where EG.5.1 predominantly circulated (**Figure 1C**). Collectively, our data suggest that BA.2.86 will spread globally and become more prevalent gradually.

**Figure 1.**
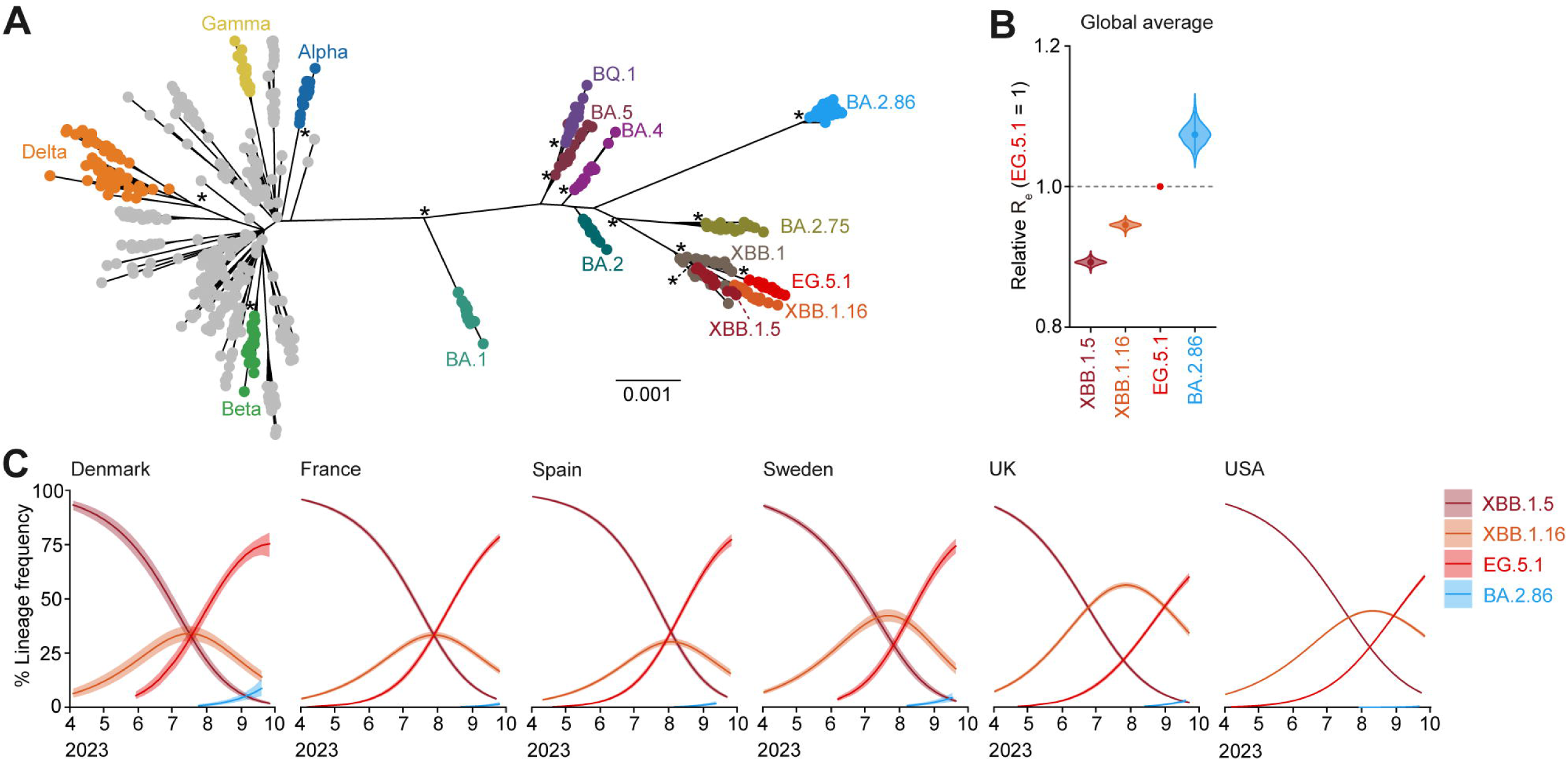
Virological features of the SARS-CoV-2 BA.2.86 (**A**) Maximum likelihood (ML) tree based on the complete genome of representative SARS-CoV-2 sequences. Twenty sequences were randomly sampled from each clade defined by Nextclade ^21^ and were included in the analysis. An asterisk represents a node with >0.95 bootstrap value. The scale bar represents genetic distance. **(B)** Estimated relative R_e_ of each viral lineages, assuming a fixed generation time of 2.1 days. The relative R_e_ of EG.5.1 is designated to 1 (horizontal dashed line). The graph includes: the posterior distribution enclosed within the 99% Bayesian confidence interval (CI; violin), the 95% CI (line), and the posterior mean (dot). The global average values estimated by a hierarchical Bayesian model ^10^ are presented. See also **Figure S1B**. **(C)** Estimated lineage dynamics of BA.2.86 and the representative SARS-CoV-2 sub-lineages in six countries where ≥20 BA.2.86 sequences were documented. The genome surveillance data from April 1, 2023, to October 2, 2023 were analyzed. The posterior mean is illustrated as the line, while the 95% Bayesian CI is shown as the ribbon. See also **Figure S1 and Table S1**.

### Virological phenotype of BA.2.86 S

To investigate the virological features of BA.2.86 S, we set out to measure the binding affinity of BA.2.86 S RBD to ACE2 receptor by the yeast display technique ^12^. We have demonstrated that the RBD of XBB.1.5 exhibits the highest binding affinity to ACE2 ^13^. Notably, the ACE2 binding affinity of BA.2.86 S RBD was comparable to that of XBB.1.5 S RBD and significantly higher than those of the S RBDs of ancestral B.1.1, XBB.1, XBB.1.16, EG.5.1 and the parental BA.2 (**Figure 2A**).

**Figure 2.**
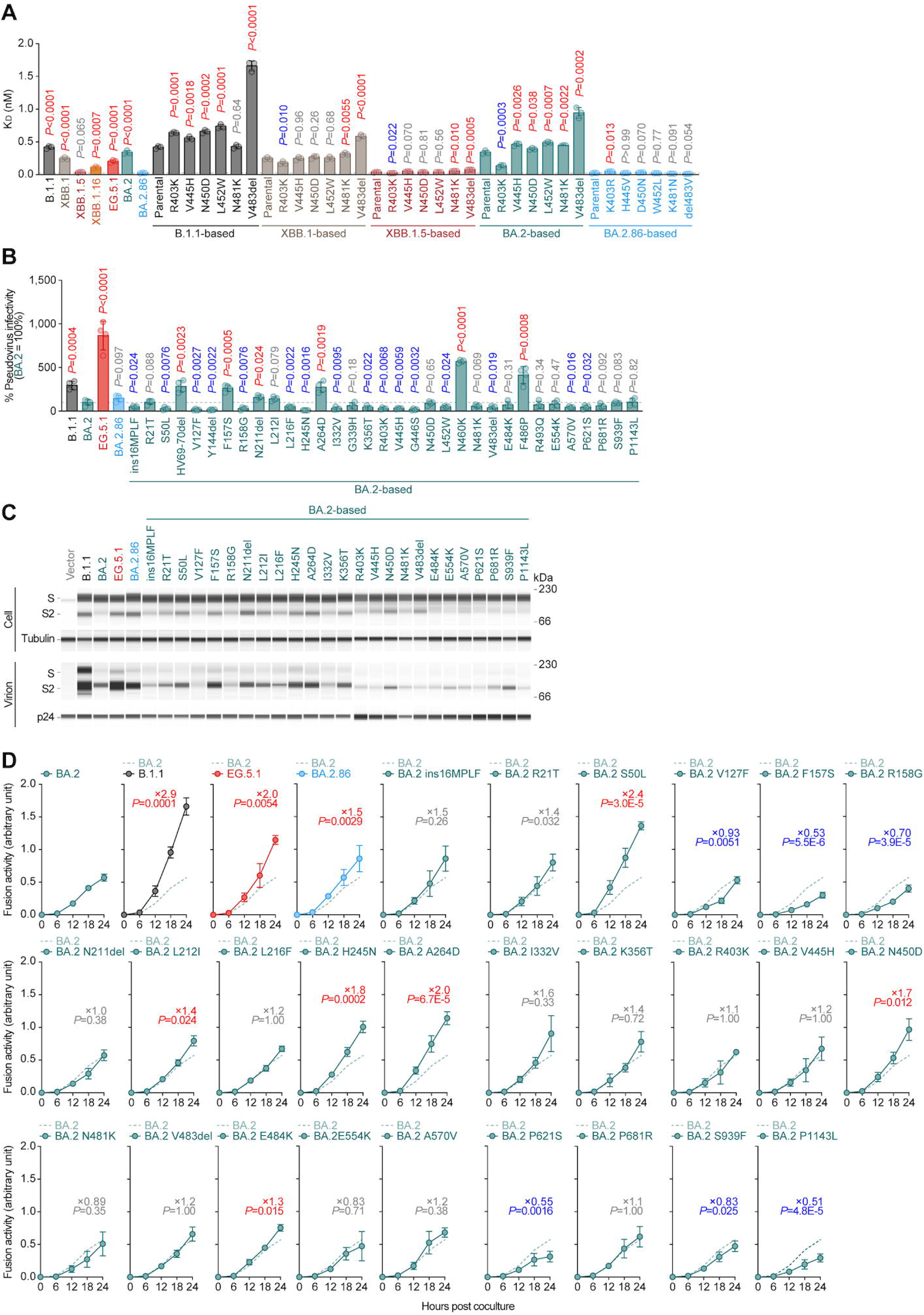
Virological phenotype of BA.2.86 S **(A)** Binding affinity of the receptor binding domain (RBD) of SARS-CoV-2 S proteins to angiotensin-converting enzyme 2 (ACE2) by yeast surface display. The dissociation constant (K_D_) value indicating the binding affinity of the RBD of the SARS-CoV-2 S protein to soluble ACE2 when expressed on yeast is shown. Each dot indicates the result of an individual replicate. The dashed horizontal lines indicate the value of BA.2. Statistically significant differences versus each parental S protein and those between BA.2 were determined by two-sided Student’s *t* tests. **(B)** Pseudovirus assay. HOS-ACE2/TMPRSS2 cells were infected with pseudoviruses bearing each S protein. The amount of input virus was normalized based on the amount of HIV-1 p24 capsid protein. The percent infectivity compared to that of the virus pseudotyped with the BA.2DS protein are shown. Assays were performed in quadruplicate. The presented data are expressed as the average ± standard deviation (SD). Each dot indicates the result of an individual replicate. The dashed horizontal lines indicate the value of BA.2. Statistically significant differences versus each parental S protein and those between BA.2 were determined by two-sided Student’s *t* tests. **(C)** Western blotting of S protein in cells and virions. Representative blots of S-expressing cells (labelled with ‘Cell’) and supernatants (labelled with ‘Virion’) are shown. Tubulin and HIV-1 p24 were used for the internal controls of ‘Cell’ and ‘Virion’, respectively. kDa, kilodalton. **(D)** S-based fusion assay in Calu-3 cells. The recorded fusion activity (arbitrary units) is shown. The dashed green line indicates the result of BA.2. The red number in each panel indicates the fold difference between BA.2 and the derivative tested at 24Dh post coculture. Assays were performed in quadruplicate. Statistically significant differences versus BA.2 across timepoints were determined by multiple regression. The familywise error rates (FWERs) calculated using the Holm method are indicated in the figures. See also **Figure S2**.

To investigate the impact of hallmark mutations in the RBD of BA.2.86 S on the binding affinity to ACE2 receptor, we generated a set of reverse mutations based on BA.2.86 S RBD. As shown in **Figure 2A**, only a substitution, the K403R, significantly increased the K_D_ value when compared to the parental BA.2.86, suggesting that the R403K substitution can lead to increased ACE2 binding affinity. The decreased K_D_ values by the R403K substitution in the S RBDs of XBB.1, XBB.1.5 and BA.2 support the observation in BA.2.86 S RBD (**Figure 2A**). However, the K_D_ value of B.1.1 R403K was significantly larger than that of parental B.1.1 (**Figure 2A**), suggesting that the effect of R403K is epistatic and the increase of ACE2 binding affinity is observed only in the case of the backbone of BA.2-related S RBD.

To test the impact of S mutations in viral infectivity, we performed an infection assay using HIV-1-based pseudovirus ^2^. The assay showed that pseudoviruses with B.1.1 or EG.5.1 S showed significantly higher infectivity than that with BA.2 S protein, but pseudovirus with BA.2.86 S protein was comparable to that with BA.2 S protein (**Figure 2B**). To test the effect of each mutation on pseudovirus infectivity, we generated a total of 33 BA.2 derivatives that bear respective mutations in BA.2.86 (**Figure 2B**). Most of the mutations did not affect BA.2 S pseudovirus infectivity or significantly decreased it (**Figure 2B**). On the other hand, consistent with our previous reports ^11,13,14^, certain mutations in the RBD, such as N460K and F486P, increased the pseudovirus infectivity (**Figure 2B**). Interestingly, three novel mutations in the N-terminal domain (NTD) of the BA.2.86 S protein, F157S, N211del, and A264D, significantly increased the pseudovirus infectivity (**Figure 2B**). To assess the association of TMPRSS2 usage with a cell entry of BA.2.86, we used HEK293-ACE2/TMPRSS2 cells and HEK293-ACE2 cells, on which endogenous surface TMPRSS2 is undetectable ^10^, as the target cells. As shown in **Figure S2A**, the impact of TMPRSS2 expression on the infectivity of BA.2.86 pseudovirus was relatively minor, suggesting that TMPRSS2 expression is not associated with a cell entry of BA.2.86.

To examine the cleavage efficiency of S protein in the cells, the cells used for pseudoviruses production were subjected to western blotting (**Figure 2C, S2B and S2C**). Interestingly, the band intensity of S2 in the cells expressing BA.2.86 S protein was higher than that of the cells expressing BA.2 S protein (**Figure 2C, S2B, S2C, S2D, and S2E**). The results from respective point mutants based on BA.2 S protein showed that multiple mutations, such as R21T, S50L, F157S, N211del, L212L, H245N, A264D, K356T, R403K, V445H, N450D, N460K, and V483del contributed to increased efficiency of S cleavage (**Figure 2C, S2B, S2C, S2D, and S2E**). In addition to the S cleavage efficacy in the cells, the level of S2 in the virions pseudotyped with BA.2.86 S protein was higher than that with BA.2 S protein (**Figure 2C, S2B and S2C**). While the levels of virion-incorporated S2 protein of respective BA.2-based point mutants were different from each other, three mutant S proteins (F157S, A264D, and N460K) with increased cleavage efficacy in the cells exhibited increased incorporation of S2 proteins in the released viral particles (**Figure 2C, S2B and S2C**).

### Fusogenicity of BA.2.86 S

We then investigated the fusogenicity of BA.2.86 S protein by the S protein-mediated membrane fusion assay in Calu-3/DSP_1-7_ cells ^15^. The surface expression level of BA.2.86 S protein was comparable to that of the parental BA.2 S protein (**Figure S2F**). Some mutations detected in BA.2.86, such as HV69-70del, Y144del, F157S, I332V, K356T, R403K, L452W, F486P, R493Q, P681R, and P1143L significantly increased the expression level of BA.2 S protein on the cell surface (**Figure S2F**).

Consistent with our previous reports ^10,16,17^, B.1.1, XBB.1.5, and EG.5.1 S proteins were significantly more fusogenic than BA.2 S protein (**Figure 2D and S2G**). Notably, the fusogenicity of BA.2.86 S protein was significantly greater than that of BA.2 S protein (**Figure 2D and S2G**). This result prompted us to determine the amino acid residues responsible for the increased fusogenicity. We found that four mutations in the NTD (S50L, L212I, H245N, A264D), and two mutations in the RBD (N450D and E484K) significantly increased the S protein fusogenicity (**Figure 2D and S2G**). Interestingly, we have demonstrated that the P681R substitution, a hallmark mutation in the Delta variant, significantly increased the fusogenicity of ancestral B.1.1-based S protein ^18^. However, the P681R substitution did not affect the fusogenicity of BA.2 S protein (**Figure 2D and Figure S2G**). Similar to the effect of R403K substitution on ACE2 binding by yeast surface display (**Figure 2A**), our results suggest that the effect of certain substitutions (e.g., R403K and P681R) on the virological feature of SARS-CoV-2 S is epistatic.

### Immune evasion of BA.2.86

We have recently reported that BA.2.86 is more resistant to XBB BTI sera than EG.5.1 ^2^. To evaluate the sensitivity of BA.2.86 to antiviral humoral immunity elicited by the breakthrough infection (BTI) with other Omicron sublineages, we performed neutralization assays using BA.2 BTI sera (n = 13) and BA.5 BTI sera (n = 17). As shown in **Figure S2H**, the 50% neutralization titer (NT_50_) of BA.2 BTI sera against BA.2.86 was significantly (43-fold) lower than those against the B.1.1 (*P* < 0.0001) as well as EG.5.1. A similar trend was observed in the BA.5 BTI sera (20-fold, *P* < 0.0001) (**Figure S2I**). These results suggest that BA.2.86 has a potent immune evasion ability from a humoral immunity induced by BA.2/BA.5 BTI. In the case of BA.2 BTI, the NT_50_ values of BA.2.86 were comparable to that of EG.5.1 (**Figure S2H**). Interestingly, however, the NT_50_ of BA.5 BTI sera against BA.2.86 showed a higher value than EG.5.1 (*P* = 0.02, **Figure S2I**), suggesting that BA.2.86 is more sensitive to BA.5 BTI sera than EG.5.1.

### Growth kinetics of clinically isolated BA.2.86 in cell cultures

To investigate the growth kinetics of BA.2.86 in *in vitro* cell cultures, clinical isolates of BA.2.86, EG.5.1, and BA.2 were inoculated into Vero cells (**Figure 3A**) and VeroE6/TMPRSS2 cells (**Figure 3B**). In Vero cells, the growth efficacy of BA.2.86 was significantly lower than that of EG.5.1 and BA.2 (**Figure 3A**). On the other hand, in VeroE6/TMPRSS2 cells, the growth kinetics of BA.2.86 and BA.2 was comparable, while BA.2.86 was less replicative than EG.5.1 (**Figure 3B**). An immunofluorescence assay at 72 h postinfection (h.p.i.) further showed that VeroE6/TMPRSS2 cells infected with BA.2.86 exhibited lower GFP intensity than EG.5.1-infected cells (**Figure 3C**). These results suggest that BA.2.86 showed a poorer replication capacity compared to EG.5.1 and BA.2.

**Figure 3.**
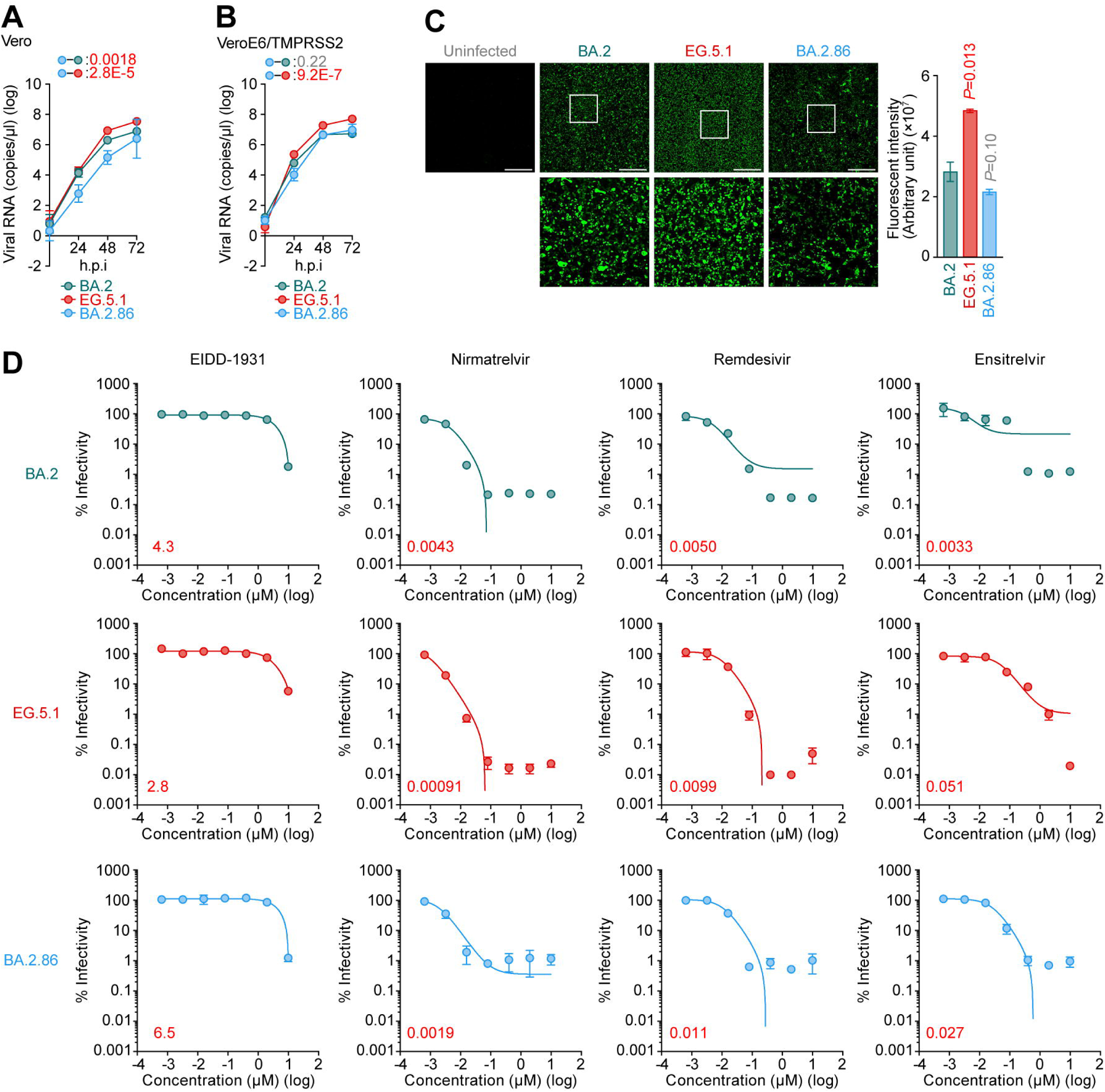
Growth kinetics of BA.2.86 and the antiviral effect of clinically available compounds against BA.2.86 (**A and B**) Growth kinetics of BA.2.86 in cell cultures. Clinical isolates of BA.2.86, EG.5.1 and BA.2 were inoculated into Vero cells (**A**) and VeroE6/TMPRSS2 cells (**B**). The copy numbers of viral RNA in the culture supernatant were routinely quantified by RT-qPCR. (**C**) Immunofluorescence staining. Infected VeroE6/TMPRSS2 cells (m.o.i. = 0.01) at 72 h.p.i. were stained with anti-SARS-CoV-2 N antibody. Higher-magnification views of the regions indicated by squares are shown. Scale bars, 1,000 μm. Left, representative panels. Higher-magnification views of the regions indicated by squares are shown at the bottom. Right, the GFP intensity of the stained cells was measured. (**D**) Effect of antiviral drugs against BA.2.86. Antiviral effects of the four drugs [EIDD-1931, Remdesivir, Ensitrelvir, and Nirmatrelvir (also known as PF-07321332)] in human iPSC-derived lung organoids. The assay of each antiviral drugs was performed in triplicate, and the 50% effective concentration (EC_50_) was calculated.

### Antiviral effect of clinically available compounds against BA.2.86

We evaluated the sensitivity of BA.2.86 to four antiviral drugs, EIDD-1931, remdesivir, ensitrelvir, and nirmatrelvir (also known as PF-07321332). Clinical isolates of BA.2 and EG.5.1 were used as controls. The viruses were inoculated into human iPSC-derived lung organoids, a physiologically relevant model, and treated with the four antiviral drugs. Nirmatrelvir showed the strongest antiviral effects, and no differences in antiviral efficacy were observed between the three variants [50% effective concentration (EC_50_) = 4.3 nM, 0.91 nM, and 1.9 nM for BA.2, EG.5.1, and BA.2.86, respectively] (**Figure 3D**). Remdesivir and ensitrelvir showed significant antiviral effects on the three isolates, while EIDD-1931 showed moderate antiviral effects on the three isolates (EC_50_ > 2 μM) (**Figure 3D**).

### Intrinsic pathogenicity of clinically isolated BA.2.86 in cell cultures

To investigate the virological features of BA.2.86 *in vivo*, clinical isolates of BA.2.86, EG.5.1, and BA.2 [2,000 50% tissue culture infectious dose (TCID_50_)] were intranasally inoculated into hamsters under anesthesia. All infected hamsters exhibited the loss of body weight (**Figure**, **A, left**). However, the loss of body weight of BA.2.86-infected hamsters was significantly less than those of the hamsters infected with EG.5.1 and BA.2 (**Figure**, **A, left**).

We then analyzed the pulmonary function of infected hamsters as reflected by two parameters, enhanced pause (Penh) and the ratio of time to peak expiratory flow relative to the total expiratory time (Rpef). Infection of EG.5.1 and BA.2 resulted in significant differences in these two respiratory parameters at 3 days postinfection (d.p.i.) (**Figure**, **A, middle** and **right**). On the other hand, these two parameters of BA.2.86-infected hamsters were constant (**Figure**, **A, middle** and **right**). These results suggest that BA.2.86 is less pathogenic in hamsters than EG.5.1 and BA.2.

To evaluate viral spread in infected hamsters, we routinely measured the viral RNA load in oral swabs and the two lung regions, lung hilum and periphery. The viral RNA load of the hamsters infected with EG.5.1 and BA.2 were comparable (**Figure**, **B**). On the other hand, the viral RNA load of BA.2.86-infected hamsters was significantly lower than those of EG.5.1- and BA.2-infected hamsters (**Figure**, **B**), suggesting that the replication efficacy of BA.2.86 *in vivo* is lower than that of EG.5.1 and BA.2.

In summary, here we elucidated the virological characteristics of BA.2.86. In our previous investigations, we observed that the S cleavage efficacy, fusogenicity, and intrinsic pathogenicity in hamsters were well correlated with each other ^10,18–20^. For instance, the Delta S protein is efficiently cleaved by furin and is highly fusogenic, and the Delta isolate is more pathogenic than ancestral SARS-CoV-2 variants ^18^. In sharp contrast, the Omicron BA.1 S protein is faintly cleaved by furin and poorly fusogenic, and the BA.1 isolate is less pathogenic than ancestral SARS-CoV-2 ^19^. Here, we showed that BA.2.86 S is more efficiently cleaved by BA.2 S, but the fusogenicity of BA.2.86 S and BA.2 S is similar. More notably, although the fusogenicity of the S protein of BA.2.86 and BA.2 were comparable, the intrinsic pathogenicity of BA.2.86 in hamsters was significantly lower than that of BA.2. This discrepancy can be explained by the replication capacity of BA.2.86. In fact, we showed that the replication kinetics of BA.2.86 is significantly lower than that of BA.2 in *in vitro* cell culture (at least in Vero cells) and *in vivo.* Therefore, our results suggest that the attenuated pathogenicity of BA.2.86 is attributed to its decreased replication capacity.

## START METHODS

● KEY RESOURCES TABLE
● RESOURCE AVAILABILITY

○ Lead Contact
○ Materials Availability
○ Data and Code Availability
● EXPERIMENTAL MODEL AND SUBJECT DETAILS

○ Ethics Statement
○ Human serum collection
○ Cell culture
● METHOD DETAILS

○ Phylogenetic analysis
○ Epidemic dynamics analysis
○ Viral genome sequencing
○ Plasmid construction
○ Yeast surface display analysis
○ Pseudovirus infection
○ Western blotting
○ SARS-CoV-2 S-based fusion assay
○ SARS-CoV-2 preparation and titration
○ SARS-CoV-2 infection
○ Immunofluorescence staining
○ RT–qPCR
○ Antiviral drug assay using SARS-CoV-2 clinical isolates and human iPSC-derived lung organoids
○ Animal experiments
○ Lung function test
● QUANTIFICATION AND STATISTICAL ANALYSIS

## Supplemental Information

Additional Supplemental Items are available upon request.

## Author Contributions

All authors reviewed and proofread the manuscript.

The Genotype to Phenotype Japan (G2P-Japan) Consortium contributed to the project administration.

## Acknowledgments

We would like to thank all members belonging to The Genotype to Phenotype Japan (G2P-Japan) Consortium. We thank Dr. Kenzo Tokunaga (National Institute for Infectious Diseases, Japan) and Dr. Jin Gohda (The University of Tokyo, Japan) for providing reagents. We thank to all members belonging to Japanese Consortium on Structural Virology (JX-Vir). We appreciate the technical assistance from The Research Support Center, Research Center for Human Disease Modeling, Kyushu University Graduate School of Medical Sciences. We gratefully acknowledge all data contributors, i.e. the Authors and their Originating laboratories responsible for obtaining the specimens, and their Submitting laboratories for generating the genetic sequence and metadata and sharing via the GISAID Initiative, on which this research is based. The super-computing resource was provided by Human Genome Center at The University of Tokyo.

This study was supported in part by AMED SCARDA Japan Initiative for World-leading Vaccine Research and Development Centers “UTOPIA” (JP223fa627001, to Kei Sato), AMED SCARDA Program on R&D of new generation vaccine including new modality application (JP223fa727002, to Kei Sato); AMED SCARDA Kyoto University Immunomonitoring Center (KIC) (JP223fa627009, to Takao Hashiguchi); AMED SCARDA Hokkaido University Institute for Vaccine Research and Development (HU-IVReD) (JP223fa627005, to Katsumi Maenaka); AMED Research Program on Emerging and Re-emerging Infectious Diseases (JP21fk0108574, to Hesham Nasser; JP21fk0108493, to Takasuke Fukuhara; JP22fk0108617 to Takasuke Fukuhara; JP22fk0108146, to Kei Sato; JP21fk0108494 to G2P-Japan Consortium, Keita Matsuno, Shinya Tanaka, Terumasa Ikeda, Takasuke Fukuhara, and Kei Sato; JP21fk0108425, to Kazuo Takayama and Kei Sato; JP21fk0108432, to Kazuo Takayama, Takasuke Fukuhara and Kei Sato; JP22fk0108534, to Takashi Irie, Terumasa Ikeda, and Kei Sato; JP22fk0108511, to Yuki Yamamoto, Akatsuki Saito, Terumasa Ikeda, Keita Matsuno, Shinya Tanaka, Kazuo Takayama, Takao Hashiguchi, Takasuke Fukuhara, and Kei Sato; JP22fk0108506, to Akatsuki Saito, Kazuo Takayama and Kei Sato); AMED Research Program on HIV/AIDS (JP23fk0410047, to Akatsuki Saito; JP23fk0410056, to Akatsuki Saito; JP23fk0410058, to Akatsuki Saito; JP22fk0410055, to Terumasa Ikeda; and JP22fk0410039, to Kei Sato); AMED Japan Program for Infectious Diseases Research and Infrastructure (JP22wm0125008 to Keita Matsuno); AMED CREST (JP21gm1610005, to Kazuo Takayama; JP22gm1610008, to Takasuke Fukuhara; JP22gm1810004, to Katsumi Maenaka); JST PRESTO (JPMJPR22R1, to Jumpei Ito); JST CREST (JPMJCR20H4, to Kei Sato; JPMJCR20H8, to Takao Hashiguchi); JSPS KAKENHI Grant-in-Aid for Scientific Research C (22K07103, to Terumasa Ikeda); JSPS KAKENHI Grant-in-Aid for Scientific Research B (21H02736, to Takasuke Fukuhara); JSPS KAKENHI Grant-in-Aid for Early-Career Scientists (22K16375, to Hesham Nasser; 20K15767, Jumpei Ito); JSPS KAKENHI grant JP20H05873 (to Katsumi Maenaka); JSPS Core-to-Core Program (A. Advanced Research Networks) (JPJSCCA20190008, to Kei Sato); JSPS Research Fellow DC2 (22J11578, to Keiya Uriu); JSPS Research Fellow DC1 (23KJ0710, to Yusuke Kosugi); JSPS Leading Initiative for Excellent Young Researchers (LEADER) (to Terumasa Ikeda); World-leading Innovative and Smart Education (WISE) Program 1801 from the Ministry of Education, Culture, Sports, Science and Technology (MEXT) (to Naganori Nao); Ministry of Health, Labour and Welfare (MHLW) under grant 23HA2010 (to Naganori Nao and Keita Matsuno); Research Support Project for Life Science and Drug Discovery [Basis for Supporting Innovative Drug Discovery and Life Science Research (BINDS)] from AMED under the Grant JP22ama121001 (to Takao Hashiguchi) and JP22ama121037 (to Katsumi Maenaka); The Cooperative Research Program (Joint Usage/Research Center program) of Institute for Life and Medical Sciences, Kyoto University (to Kei Sato and Katsumi Maenaka); International Joint Research Project of the Institute of Medical Science, the University of Tokyo (to Akatsuki Saito, Terumasa Ikeda, Jiri Zahradnik, and Takasuke Fukuhara); The Tokyo Biochemical Research Foundation (to Kei Sato); Takeda Science Foundation (to Terumasa Ikeda and Katsumi Maenaka); Mochida Memorial Foundation for Medical and Pharmaceutical Research (to Terumasa Ikeda); The Naito Foundation (to Terumasa Ikeda); Mitsubishi Foundation (to Kei Sato); Japanese Government MEXT Scholarship-Research Category (220235, Jarel Elgin Tolentino) and the project of National Institute of Virology and Bacteriology, Programme EXCELES, funded by the European Union, Next Generation EU (LX22NPO5103, to Jiri Zahradnik).

## Consortia

The Genotype to Phenotype Japan (G2P-Japan) Consortium

## STARTMETHODS

### KEY RESOURCES TABLE

### RESOURCE AVAILABILITY

#### Lead Contact

Further information and requests for resources and reagents should be directed to and will be fulfilled by the Lead Contact, Kei Sato.

#### Materials Availability

All unique reagents generated in this study are listed in the Key Resources Table and available from the Lead Contact with a completed Materials Transfer Agreement.

#### Data and Software Availability

All databases/datasets used in this study are available from GenBank database (https://www.ncbi.nlm.nih.gov/genbank/) the GISAID database (https://www.gisaid.org; EPI_SET_230919bh; EPI_SET_231030mc). Computational codes used in this study are available on the GitHub repository (https://github.com/TheSatoLab/BA.2.86_full).

### EXPERIMENTAL MODEL AND SUBJECT DETAILS

#### Ethics statement

All experiments with hamsters were performed in accordance with the Science Council of Japan’s Guidelines for the Proper Conduct of Animal Experiments. The protocols were approved by the Institutional Animal Care and Use Committee of National University Corporation Hokkaido University (approval ID: 20-0123 and 20-0060). All protocols involving specimens from human subjects recruited at Interpark Kuramochi Clinic were reviewed and approved by the Institutional Review Board of Interpark Kuramochi Clinic (approval ID: G2021-004). All human subjects provided written informed consent. All protocols for the use of human specimens were reviewed and approved by the Institutional Review Boards of The Institute of Medical Science, The University of Tokyo (approval IDs: 2021-1-0416 and 2021-18-0617).

#### Human serum collection

Convalescent sera were collected from fully vaccinated individuals who had been infected with BA.2 (9 2-dose vaccinated and 4 3-dose vaccinated; 11–61 days after testing. n=13 in total; average age: 45 years, range: 24–82 years, 62% male) (**Figure S2H**), and fully vaccinated individuals who had been infected with BA.5 (1 2-dose vaccinated, 15 3-dose vaccinated and 1 4-dose vaccinated; 10–23 days after testing. n=17 in total; average age: 52 years, range: 25–73 years, 53% male) (**Figure S2I**). The SARS-CoV-2 variants were identified as previously described ^10,20^. Sera were inactivated at 56°C for 30 minutes and stored at –80°C until use. The details of the convalescent sera are summarized in **Table S2**.

#### Cell culture

HEK293T cells (a human embryonic kidney cell line; ATCC, CRL-3216), HEK293 cells (a human embryonic kidney cell line; ATCC, CRL-1573), LentiX-293T (a derivative of HEK293T cells for superior lentivirus packaging; TaKaRa, Cat# 632180) and HOS-ACE2/TMPRSS2 cells (HOS cells stably expressing human ACE2 and TMPRSS2) ^22,23^ were maintained in DMEM (high glucose) (Sigma-Aldrich, Cat# 6429-500ML) containing 10% fetal bovine serum (FBS, Sigma-Aldrich Cat# 172012-500ML) and 1% penicillin–streptomycin (PS) (Sigma-Aldrich, Cat# P4333-100ML). HEK293-ACE2 cells (HEK293 cells stably expressing human ACE2) ^24^ were maintained in DMEM (high glucose) containing 10% FBS, 1 µg/ml puromycin (InvivoGen, Cat# ant-pr-1) and 1% PS. HEK293-ACE2/TMPRSS2 cells (HEK293 cells stably expressing human ACE2 and TMPRSS2) ^24^ were maintained in DMEM (high glucose) containing 10% FBS, 1 µg/ml puromycin, 200 µg/ml hygromycin (Nacalai Tesque, Cat# 09287-84) and 1% PS. Calu-3/DSP_1-7_ cells (Calu-3 cells stably expressing DSP_1-7_) ^25^ were maintained in EMEM (Wako, Cat# 056-08385) containing 20% FBS and 1% PS. Vero cells [an African green monkey (Chlorocebus sabaeus) kidney cell line; JCRB Cell Bank, JCRB0111] were maintained in Eagle’s minimum essential medium (EMEM) (Sigma-Aldrich, Cat#M4655-500ML) containing 10% FBS and 1% PS. VeroE6/TMPRSS2 cells (VeroE6 cells stably expressing human TMPRSS2; JCRB Cell Bank, JCRB1819) ^26^ were maintained in DMEM (low glucose) (Wako, Cat#041-29775) containing 10% FBS, G418 (1 mg/ml; Nacalai Tesque, Cat#G8168-10ML) and 1% PS.

### METHOD DETAILS

#### Phylogenetic analysis

A total of 15,991,922 SARS-CoV-2 genome sequences and their metadata were downloaded from the GISAID database with a released date of September 14, 2023 (https://www.gisaid.org/). To prepare dataset for lineages other than BA.2.86, the dataset was then filtered based on the following criteria: (i) retained only distinct Accession IDs, (ii) host labeled as ‘Human’, (iii) the collection date recorded, (iv) the PANGO lineage column should not be empty, none or unassigned, and (v) retained sequences with less than 1% proportion of ambiguous bases. We assigned Nextclade clade information to individual viral sequences using the Nextclade v2.14.1 CLI workflow (https://clades.nextstrain.org/). Subsequently, we randomly sampled 20 sequences from each Nextclade clade. To prepare dataset for BA.2.86 (including BA.2.86.1), we extracted sequences in which PANGO lineage is BA.2.86 or BA.2.86.1 from the GISAID metadata. Subsequently, we applied the same filtering criteria as mentioned above (i-iv) and additionally set the threshold for ambiguous bases below 3%. We set this relaxed threshold for BA.2.86 because most of BA.2.86 sequences have a large undetermined regions just before S gene due to the presence of mutations in the primer site. After the filtering, 89 sequences of BA.2.86 were included in the final dataset.

To construct the phylogenetic tree, viral genome sequences (EPI SET ID: EPI_SET_230919bh) were mapped and aligned to the reference sequence of Wuhan-Hu-1 (GenBank accession number: NC_045512.2) through minimap v2.24 ^27^, and the resulting sam format file was converted to fasta format using gofasta v1.2.0 ^28^. During this conversion, the alignment sites corresponding to 1–265and 29674–9903 positions on the reference genome were masked, typically converted to ‘NNN’. Alignment sites with more than 10% of sequences containing gaps or uncertain nucleotides were subjected to trimming using trimAl v1.2 ^29^. Phylogenetic tree construction was accomplished via the three-step protocol: (i) the initial tree was constructed, (ii) the external branch lengths of the initial tree were filtered using Grubb’s test and the *p value* threshold was set to 1.0E-5 enabling those tips with longer external branch to be removed, (iii) the final tree was constructed with the similar parameter as the initial tree ^14^. A maximum likelihood (ML) phylogenetic tree of the genome was inferred by IQTree v2.2.2.6 with the GTR nucleotide substitution model ^30^. The node support value was computed by 1000 bootstrap iterations. The visualization of the final tree was generated in R v4.3.1 using the ggtree package ^31^.

#### Epidemic dynamics analysis

To estimate the global average and country-specific R_e_ values of SARS-CoV-2 lineages, we analyzed the GISAID genome surveillance data spanning from April 1, 2023 to October 2, 2023. Genomic and surveillance data of 16,063,834 sequences with a released date of October 2, 2023, were acquired from the GISAID database (https://www.gisaid.org/). We excluded the sequence records with the following features: i) a lack of collection date information; ii) sampling in animals other than humans; iii) sampling by quarantine; or iv) without the PANGO lineage information. We then allocated Nextclade clade information to individual viral sequences using the Nextclade v2.14.1 CLI workflow (https://clades.nextstrain.org/). For the definition of lineages other than BA.2.86, we used the Nextclade clade classification: 23A (XBB.1.5), 23B (XBB.1.16), and 23F (EG.5.1). Since BA.2.86 (including BA.2.86.1) has not been annotated in the Nextclade clade, we instead used PANGO lineage classification assigned by Nextclade for these lineages. BA.2.86 sublineages (e.g., BA.2.86.1) are summarized as BA.2.86. We then analyzed the datasets of the countries with ≥20 available BA.2.86 sequences: Denmark, France, Spain, Sweden, UK, and USA (analyzed dataset: EPI_SET_231030mc). Subsequently, we counted the daily frequency of each viral lineage in each country and fit a Bayesian hierarchical multinomial logistic model ^10,11^ to the lineage frequency data to estimate the global average and country-specific R_e_ of the lineages. The relative R_e_ of each viral lineage l in each county s (r_lS_) was calculated according to the country-specific slope parameter, β_lS_, as r_lS_ = exp(γβ_lS_) where y is the average viral generation time (2.1 days)(http://sonorouschocolate.com/covid19/index.php?title=Estimating_Gener <u>ation_Time_Of_Omicron</u>). Similarly, the global average relative R_e_ of each viral lineage was calculated according to the global average slope parameter, β_l_, as r_l_ = exp(γβ_l_). For parameter estimation, the intercept and slope parameters of the EG.5.1 were set at 0. As a result, the relative R_e_ of EG.5.1 was fixed at 1, and the R_e_ of other viral lineages were estimated relative to that of EG.5.1. Parameter estimation was conducted via the MCMC method implemented in CmdStan v2.33 (https://mc-stan.org) with CmdStanr v0.6.1 (https://mc-stan.org/cmdstanr/). Four separate MCMC chains were executed, consisting of 1,000 steps as the warmup iterations, and 2,000 steps as the sampling iterations. We verified the successful convergence of our MCMC runs by assuring that all the estimated parameters had showed <1.01 R-hat convergence diagnostic values and >200 effective sampling size values. Information on the estimated parameters is summarized in **Table S1**.

#### Viral genome sequencing

Viral genome sequencing was performed as previously described ^20^. Briefly, the virus sequences were verified by viral RNA-sequencing analysis. Viral RNA was extracted using a QIAamp viral RNA mini kit (Qiagen, Cat# 52906). The sequencing library employed for total RNA sequencing was prepared using the NEBNext Ultra RNA Library Prep Kit for Illumina (New England Biolabs, Cat# E7530). Paired-end 76-bp sequencing was performed using a MiSeq system (Illumina) with MiSeq reagent kit v3 (Illumina, Cat# MS-102-3001). Sequencing reads were trimmed using fastp v0.21.0 ^32^ and subsequently mapped to the viral genome sequences of a lineage B isolate (strain Wuhan-Hu-1; GenBank accession number: NC_045512.2) ^26^ using BWA-MEM v0.7.17 ^33^. Variant calling, filtering, and annotation were performed using SAMtools v1.9 ^34^ and snpEff v5.0e ^35^.

#### Plasmid construction

Plasmids expressing the codon-optimized SARS-CoV-2 S proteins of B.1.1 (the parental D614G-bearing variant), BA.2, EG.5.1, and BA.2.86 were prepared in our previous studies ^2,17,20^. Plasmids expressing the codon-optimized S proteins of BA.2.86 and BA.2 S-based derivatives were generated by site-directed overlap extension PCR using the primers listed in **Table S3**. The resulting PCR fragment was digested with KpnI (New England Biolabs, Cat# R0142S) and NotI (New England Biolabs, Cat# R1089S) and inserted into the corresponding site of the pCAGGS vector ^36^. Nucleotide sequences were determined by DNA sequencing services (Eurofins), and the sequence data were analyzed by Sequencher v5.1 software (Gene Codes Corporation). Nucleotide sequences were determined by DNA sequencing services (Eurofins), and the sequence data were analyzed by Sequencher v5.1 software (Gene Codes Corporation).

#### Yeast surface display analysis

Utilizing yeast surface display (**Figure 2A**), we conducted an analysis of the interaction between selected RBD variants and mACE2, following established protocols ^11–14,17,37–39^. The pJYDC plasmids bearing SARS-CoV-2_RBD-WT, BA2 XBB, XBB.1.5, XBB.1.16 and EG.5.1 variants were used in our previous research ^2,10,13,17,38–40^. The gene for RBD-BA.2.86 with *S. cerevisiae* codon usage was obtained from Twist Biosciences. The mutations in RBDs were incorporated by restriction-free cloning. All PCR reactions were conducted using the KAPA HiFi HotStart ReadyMix kit (Roche, Cat# KK2601) and the pJYDC1 plasmid (Addgene, Cat# 162458), as previously outlined ^2,10,13,17,38–40^. A detailed list of the primers used can be found in **Table S3**. Verified plasmids were transformed into yeast Saccharomyces cerevisiae strain EBY100 (ATCC, MYA-4941) through electroporation and selected on SD-Trp selection plates. Yeast colonies were grown for 24 h in the liquid culture (SDCAA, 30°C, 220 rpm) and the yeast expression proceeded for 48 h at 20°C in 1/9 media. Expressed yeasts were washed with PBS supplemented with bovine serum albumin at a concentration of 1 g/l (PBSB). The cells were then exposed to a range of mACE2 concentrations (4 pM to 10 nM, in a dilution series with a factor of 2) and 20 nM bilirubin (Sigma-Aldrich, Cat# 14370-1G), washed with PBSB and the recorded data included RBD expression and ACE2 signal, captured using automated acquisition from 96-well plates by the FACS CytoFLEX Flow Cytometer (Beckman Coulter). Background binding signals were subtracted, and fluorescence spill of eUnaG2 signals into the red channel was compensated. Subsequently, the data were fitted to a standard noncooperative Hill equation through nonlinear least-squares regression, utilizing Python v3.7 (https://www.python.org) as previously detailed ^2,10,13,17,38–40^.

#### Pseudovirus infection

Pseudovirus infection (Figures 2B and S2A) was performed as previously described ^19,23,24,41–44^. Briefly, lentivirus (HIV-1)-based, luciferase-expressing reporter viruses were pseudotyped with the SARS-CoV-2 S protein. One prior day of transfection, the LentiX-293T or HEK293T cells were seeded at a density of 2 × 10^6^ cells. The LentiX-293T or HEK293T cells were cotransfected with 1 μg psPAX2-IN/HiBiT (a packaging plasmid encoding the HiBiT-tag-fused integrase ^45^, 1 μg pWPI-Luc2 (a reporter plasmid encoding a firefly luciferase gene ^45^ and 500 ng plasmids expressing parental S protein or its derivatives using TransIT-293 transfection reagent (Mirus, Cat# MIR2704) or TransIT-LT1 (Takara, Cat# MIR2300) according to the manufacturer’s protocol. Two days posttransfection, the culture supernatants were harvested and filtrated. The amount of produced pseudovirus particles was quantified by the HiBiT assay using the Nano Glo HiBiT lytic detection system (Promega, Cat# N3040) as previously described ^45^. In this system, HiBiT peptide is produced with HIV-1 integrase and forms NanoLuc luciferase with LgBiT, which is supplemented with substrates. In each pseudovirus particle, the detected HiBiT value is correlated with the amount of the pseudovirus capsid protein, HIV-1 p24 protein ^45^. Therefore, we calculated the amount of HIV-1 p24 capsid protein based on the HiBiT value measured, according to the previous paper ^45^. To measure viral infectivity, the same amount of pseudovirus normalized with the HIV-1 p24 capsid protein was inoculated into HOS-ACE2/TMPRSS2 cells, HEK293-ACE2, and HEk293-ACE2/TMPRSS2 cells. At two days postinfection, the infected cells were lysed with a Bright-Glo luciferase assay system (Promega, Cat# E2620), and the luminescent signal produced by firefly luciferase reaction was measured using a GloMax explorer multimode microplate reader 3500 (Promega) or CentroXS3 LB960 (Berthold Technologies). The pseudoviruses were stored at –80°C until use. To analyze the effect of TMPRSS2 for pseudovirus infectivity (Figure S2A), the fold change of the values of HEK293-ACE2/TMPRSS2 to HEK293-ACE2 was calculated.

#### Western blotting

As previously described, sample preparation for western blotting was performed with minor modifications ^15,46^. For western blotting, HEK293T cells (2 × 10^6^ cells) were cotransfected with 2 μg of psPAX2-IN/HiBiT, 2 μg of pWPI-Luc2, and 1 μg of plasmids expressing SARS-CoV-2 S using TransIT-LT1 according to the manufacturer’s protocol. At 2 days posttransfection, cell culture supernatants were collected, filtered, and subjected to ultracentrifugation using 20% sucrose (22,000 × *g*, 4°C, 2 hours). Then, virions were dissolved in phosphate-buffered saline (PBS). To quantify HIV-1 p24 antigen in the pseudovirus, the amount of pseudoviruses in the cell culture supernatant was quantified by the HiBiT assay using a Nano Glo HiBiT lytic detection system (Promega, Cat# N3040). After normalization with HiBiT value, the samples were diluted with 2 × SDS sample buffer [100 mM Tris-HCl (pH6.8), 4% SDS, 12% β-mercaptoethanol, 20% glycerol, 0.05% bromophenol blue] and boiled for 5–10Dminutes at 100°C. For cell lysate preparation, the transfected cells were detached, washed twice with PBS, and lysed in lysis buffer [25mM HEPES (pH7.2), 20% glycerol, 125 mM NaCl, 1% Nonidet P40 substitute (Nacalai Tesque, Cat# 18558-54), protease inhibitor cocktail (Nacalai Tesque, Cat# 03969-21)]. Quantification of total protein in the cell lysates was done by protein assay dye (Bio-Rad, Cat# 5000006) according to manufacturer’s instruction. Then, cell lysates were diluted with 2 × SDS sample buffer and boiled for 5–10Dminutes. After cooling down, viral (pseudovirus) and cell lysates were mixed with diluted sample buffer (proteinsimple, Cat# 99351). Then, 5 × Fluorescent Master mix (proteinsimple, Cat# PS-ST01EZ-8) was added at a ratio of 4:1. Simple Western System, Abby (proteinsimple) was used for protein analysis. For protein detection, the following antibodies were used: rabbit anti-SARS-CoV-2 S (Novus Biologicals, Cat# NB100-56578, viral lysate; 1:40, cell lysate; 1:40). mouse anti-HIV-1 p24 monoclonal antibody (HIV Reagent Program, ARP-3537, 1:500), mouse anti-α tubulin monoclonal antibody (Sigma-Aldrich, Cat# T5168, 1:100), anti-rabbit secondary antibody (proteinsimple, Cat# 042-206), and anti-mouse secondary antibody (proteinsimple, Cat# 042-205). Bands were visualized and analyzed using Compass for Simple Western v6.1.0 (proteinsimple).

#### SARS-CoV-2 S-based fusion assay

A SARS-CoV-2 S-based fusion assay (**Figures 2D, S2F and S2G**) was performed as previously described ^10,11,14–17,19,20,24,39,47^. Briefly, on day 1, effector cells (i.e., S-expressing cells) and target cells (Calu-3/DSP_1-7_ cells) were prepared at a density of 0.6–0.8 × 10^6^ cells in a 6-well plate. On day 2, for the preparation of effector cells, HEK293 cells were cotransfected with the S expression plasmids (400 ng) and pDSP_8-11_ ^48^ (400 ng) using TransIT-LT1 (Takara, Cat# MIR2300). On day 3 (24 hours posttransfection), 16,000 effector cells were detached and reseeded into a 96-well black plate (PerkinElmer, Cat# 6005225), and target cells were reseeded at a density of 1,000,000 cells/2 ml/well in 6-well plates. On day 4 (48 hours posttransfection), target cells were incubated with EnduRen live cell substrate (Promega, Cat# E6481) for 3 hours and then detached, and 32,000 target cells were added to a 96-well plate with effector cells. *Renilla* luciferase activity was measured at the indicated time points using Centro XS3 LB960 (Berthhold Technologies). For measurement of the surface expression level of the S protein, effector cells were stained with rabbit anti-SARS-CoV-2 S S1/S2 polyclonal antibody (Thermo Fisher Scientific, Cat# PA5-112048, 1:100). Normal rabbit IgG (Southern Biotech, Cat# 0111-01, 1:100) was used as a negative control, and APC-conjugated goat anti-rabbit IgG polyclonal antibody (Jackson ImmunoResearch, Cat# 111-136-144, 1:50) was used as a secondary antibody. The surface expression level of S proteins (Figure S2F) was measured using CytoFLEX Flow Cytometer (Beckman Coulter) and the data were analyzed using FlowJo software v10.7.1 (BD Biosciences). For calculation of fusion activity, *Renilla* luciferase activity was normalized to the mean fluorescence intensity (MFI) of surface S proteins. The normalized value (i.e., *Renilla* luciferase activity per the surface S MFI) is shown as fusion activity.

#### SARS-CoV-2 preparation and titration

The working virus stocks of SARS-CoV-2 were prepared and titrated as previously described ^10,11,14,17–20,24,39,43^. In this study, clinical isolates of BA.2.86 (strain TKYnat15020; GISAID ID: EPI_ISL_18233521), EG.5.1 (strain KU2023071028; GISAID ID: EPI_ISL_18072016) ^17^, and BA.2 (strain TY40-385; PANGO lineage BA.2, GISAID ID: EPI_ISL_9595859) ^20^ were used. The working virus stocks of BA.2 and EG.5.1 were prepared in our previous studies ^17,20^. To prepare the working virus stock of BA.2.86, 100 μl of the seed virus was inoculated into VeroE6/TMPRSS2 cells (1,000,000 cells in a one-well of 6-well plate). After 1 h absorption, the cells were cultured with DMEM (low glucose) (Fujiflim Wako, Cat# 041-29775) containing 2% FBS and 1% PS. At 3 d.p.i., the culture medium was harvested and then, subjected to inoculation into the naïve Vero/E6/TMPRS2 cells (10,000,000 cells in a 100-mm culture dish). After 84 h.p.i, the culture medium was harvested and centrifuged. The resultant supernatants were collected as the working virus stock.

The titer of the prepared working virus was measured as the 50% tissue culture infectious dose (TCID_50_). Briefly, one day before infection, VeroE6/TMPRSS2 cells (10,000 cells) were seeded into a 96-well plate. Serially diluted virus stocks were inoculated into the cells and incubated at 37°C for 4 d. The cells were observed under a microscope to judge the CPE appearance. The value of TCID_50_/ml was calculated with the Reed–Muench method ^49^.

For verification of the sequences of SARS-CoV-2 working viruses, viral RNA was extracted from the working viruses using a QIAamp viral RNA mini kit (Qiagen, Cat# 52906) and viral genome sequences were analyzed as described above (see “Viral genome sequencing” section). Information on the unexpected substitutions detected is summarized in **Table S4** and the raw data are deposited in the GitHub repository (https://github.com/TheSatoLab/BA.2.86_full1).

#### SARS-CoV-2 infection

One day before infection, Vero cells (10,000 cells) and VeroE6/TMPRSS2 cells (10,000 cells) were seeded into a 96-well plate. SARS-CoV-2 [100 TCID_50_ for Vero cells (Figure 3A) and VeroE6/TMPRSS2 cells (Figure 3B)] was inoculated and incubated at 37°C for 1 h. The infected cells were washed, and 180 µl culture medium was added. The culture supernatant (10 µl) was harvested at the indicated timepoints and used for RT–qPCR to quantify the viral RNA copy number (see “RT–qPCR” section below).

#### Immunofluorescence staining

Immunofluorescence staining (**Figure 3C**) was performed as previously described ^18,19^. In brief, one day before infection, VeroE6/TMPRSS2 cells (10,000 cells) were seeded into 96-well, glass bottom, black plates and infected with SARS-CoV-2 (100 TCID_50_). At 72 h.p.i., the cells were fixed with 4% para-formaldehyde in phosphate-buffered saline (PBS) (Nacalai Tesque, 09154-85) for 1 h at 4 °C. The fixed cells were permeabilized with 0.2% Triton X-100 in PBS for 1 h and blocked with 10% FBS in PBS for 1 h at 4 °C. The fixed cells were then stained using rabbit anti-SARS-CoV-2 N poly-clonal antibody (GeneTex, GTX135570, 1:1,000) for 1 h. After washing three times with PBS, cells were incubated with an Alexa 488-conjugated anti-rabbit IgG antibody (Thermo Fisher Scientific, A-11008, 1:1,000) for 1 h. Fluorescence microscopy was performed on an All-in-One Fluorescence Microscope BZ-X800 (Keyence). Captured images were reconstructed and the fluorescent intensity was measured by using a BZ-X800 Analyzer software (Keyence).

#### RT–qPCR

RT–qPCR was performed as previously described ^10,11,14,16–20,24,39,50^. Briefly, 5 μl culture supernatant was mixed with 5 μl 2 × RNA lysis buffer [2% Triton X-100 (Nacalai Tesque, Cat# 35501-15), 50 mM KCl, 100 mM Tris-HCl (pH 7.4), 40% glycerol, 0.8 U/μl recombinant RNase inhibitor (Takara, Cat# 2313B)] and incubated at room temperature for 10 m. RNase-free water (90 μl) was added, and the diluted sample (2.5 μl) was used as the template for real-time RT-PCR performed according to the manufacturer’s protocol using One Step TB Green PrimeScript PLUS RT-PCR kit (Takara, Cat# RR096A) and the following primers: Forward *N*, 5’-AGC CTC TTC TCG TTC CTC ATC AC-3’; and Reverse *N*, 5’-CCG CCA TTG CCA GCC ATT C-3’. The viral RNA copy number was standardized with a SARS-CoV-2 direct detection RT-qPCR kit (Takara, Cat# RC300A). Fluorescent signals were acquired using QuantStudio 3 Real-Time PCR system (Thermo Fisher Scientific), CFX Connect Real-Time PCR Detection system (Bio-Rad), Eco Real-Time PCR System (Illumina), qTOWER3 G Real-Time System (Analytik Jena) or 7500 Real-Time PCR System (Thermo Fisher Scientific).

#### Antiviral drug assay using SARS-CoV-2 clinical isolates and human iPSC-derived lung organoids

Antiviral drug assay (**Figure 3D**) was performed as previously described ^43^. Human iPSC-derived lung organoids were generated as previously described ^51^. The human iPSC-derived lung organoids were infected with either BA.2, EG.5.1, or BA.2.86 isolate (100 TCID50) at 37 °C for 2 h. The cells were washed with DMEM and cultured in DMEM supplemented with 10%DFCS, 1% PS and the serially diluted EIDD-1931 (an active metabolite of Molnupiravir; Cell Signalling Technology, Cat# 81178S), Remdesivir (Clinisciences, Cat# A17170), Ensitrelvir (MedChemExpress, Cat# HY-143216), or Nirmatrelvir (PF-07321332; MedChemExpress, Cat# HY-138687). At 72 h after the infection, the culture supernatants were collected, and viral RNA was quantified using RT–qPCR (see “RT-qPCR” section above). The assay of each compound was performed in triplicate, and the 50% effective concentration (EC_50_) was calculated using Prism 9 software v9.1.1 (GraphPad Software).

#### Animal experiments

Animal experiments (**Figure 4**) were performed as previously described^10,11,14,16–20,39,50^. Syrian hamsters (male, 4 weeks old) were purchased from Japan SLC Inc. (Shizuoka, Japan). Baseline body weights were measured before infection. For the virus infection experiments, hamsters were anaesthetized by intramuscular injection of a mixture of either 0.15 mg/kg medetomidine hydrochloride (Domitor^®^, Nippon Zenyaku Kogyo), 2.0 mg/kg midazolam (FUJIFILM Wako Chemicals, Cat# 135-13791) and 2.5 mg/kg butorphanol (Vetorphale^®^, Meiji Seika Pharma), or 0.15 mg/kg medetomidine hydrochloride, 2.0 mg/kg alphaxaone (Alfaxan^®^, Jurox) and 2.5 mg/kg butorphanol. Clinical isolates of SARS-CoV-2 (BA.2.86, BA.2, and EG.5.1) (2,000 TCID_50_ in 100 µl), or medium (100 µl) were intranasally inoculated under anesthesia. Oral swabs were collected at 2 and 5 d.p.i. Body weight, enhanced pause (Penh) and the ratio of time to peak expiratory follow relative to the total expiratory time (Rpef) were routinely monitored at indicated timepoints (see “Lung function test” section below). Respiratory organs were anatomically collected at 1, 3 and 5 d.p.i (for lung) or 1 d.p.i. (for trachea). Viral RNA load in the respiratory tissues and oral swab were determined by RT–qPCR. The respiratory tissues were also used for histopathological and IHC analyses (see “H&E staining” and “IHC” sections below). Sera of infected hamsters were collected at 16 d.p.i. using cardiac puncture under anesthesia with isoflurane and used for neutralization assay (see “Neutralization assay” above).

**Figure 4.**
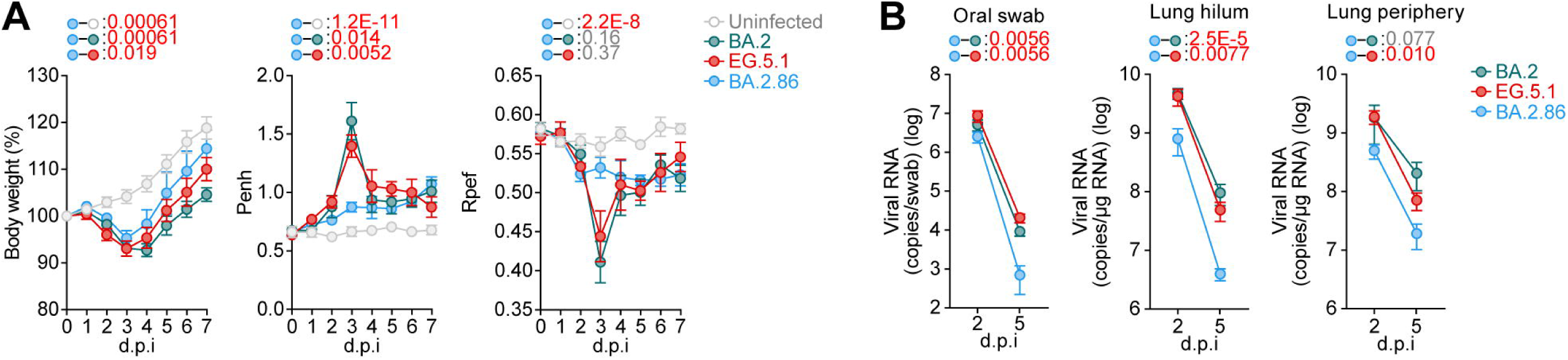
Virological features of BA.2.86 *in vivo* Syrian hamsters were intranasally inoculated with BA.2.86, EG.5.1 and BA.2. Six hamsters of the same age were intranasally inoculated with saline (uninfected). **(A)** Time-course analysis. Six hamsters per group were used to routinely measure body weight (**A**, left), Penh (**A**, middle), and Rpef (**A**, right). **(B)** Viral RNA load. Four hamsters per group were euthanized at 2 and 5 d.p.i. and quantified viral RNA load in oral swab (**B**, left), lung hilum (**B**, middle), and lung periphery (**B**, right) by RT–qPCR. In **A,B**, data are presented as the average ± SEM. In **A,B**, statistically significant differences between EG.5.1, EG5.1.1 and other variants across timepoints were determined by multiple regression. In **A**, the 0 d.p.i. data were excluded from the analyses. The FWERs calculated using the Holm method are indicated in the figures.

#### Lung function test

Lung function test (**Figure 4A**) was performed every day as previously described^10,11,14,16–20,39^. Respiratory parameters (Penh and Rpef) were measured by using a whole-body plethysmography system (DSI) according to the manufacturer’s instructions. In brief, a hamster was placed in an unrestrained plethysmography chamber and allowed to acclimatize for 30 seconds, then, data were acquired over a 2.5-minute period by using FinePointe Station and Review softwares v2.9.2.12849 (STARR).

### QUANTIFICATION AND STATISTICAL ANALYSIS

Statistical significance was tested using a two-sided Mann–Whitney *U* test, a two-sided Student’s *t* test, a two-sided Welch’s *t* test, or a two-sided paired *t-*test unless otherwise noted. The tests above were performed using Prism 9 software v9.1.1 (GraphPad Software).

In the time-course experiments (**Figures 2D, 3A, 3B, 4A, 4B, and S2G**), a multiple regression analysis including experimental conditions (i.e., the types of infected viruses) as explanatory variables and timepoints as qualitative control variables was performed to evaluate the difference between experimental conditions thorough all timepoints. The initial time point was removed from the analysis. The *P* value was calculated by a two-sided Wald test. Subsequently, familywise error rates (FWERs) were calculated by the Holm method. These analyses were performed in R v4.1.2 (https://www.r-project.org/).

In Figure 3C, photographs shown are the representatives of 57 fields of view taken for each sample of at least two independent experiments.

**Table S1.Estimated relative Re values of viral lineages by a hierarchical Bayesian model, related to Figure 1**

**Table S2. Human sera used in this study, related to Figure S2**

**Table S3. Primers used in this study, related to Figure 2**

**Table S4. Summary of unexpected amino acid mutations detected in the working virus stocks, related to Figure 3**

**Figure S1. Epidemic dynamics of BA.2.86, related to Figure 1**

**(A)** Mutation frequencies in BA.2.86 (including BA.2.86.1) and other SARS-CoV-2 lineages BA.2, BA.2.75, XBB.1, and XBB.1.5. Mutations with a frequency of >0.5 values in at least one but not all subvariants of interest are demonstrated.

**(B)** Estimated relative R_e_ of each viral lineage in each country. The graph includes: the posterior distribution enclosed within the 99% Bayesian confidence interval (CI; violin), the 95% CI (line), and the posterior mean (dot). The country-specific values estimated by a hierarchical Bayesian model ^10^ are shown.

**Figure S2. Virological features of BA.2.86 S, related to Figure 2**

**(A)** TMPRSS2 usage. Fold increase in pseudovirus infectivity based on TMPRSS2 expression was tested by infecting HEK293-ACE2 cells and HEK293-ACE2/TMPRSS2 cells with pseudoviruses bearing each S protein. Assays were performed in quadruplicate. The presented data are expressed as the average ± standard deviation (SD). Each dot indicates the result of an individual replicate. The dashed horizontal lines indicate the value of BA.2. Statistically significant differences versus each parental S protein and those between BA.2 were determined by two-sided Student’s *t* tests.

**(B and C)** Western blotting of S protein in cells and virions. Representative blots of S-expressing cells (labelled with ‘Cell’) and supernatants (labelled with ‘Virion’) are shown. Tubulin and HIV-1 p24 were used for the internal controls of ‘Cell’ and ‘Virion’, respectively. kDa, kilodalton.

**(D and E)** S cleavage efficiency. Each bar indicates S2/(S + S2) ratio relative to BA.2 with the mean ± SD from 4 independent experiments. The dashed horizontal lines indicate the value of BA.2. Statistically significant differences versus each parental S protein and those between BA.2 were determined by two-sided Student’s *t* tests.

**(F)** S protein expression on the cell surface. Mean fluorescence intensity (MFI) of surface S protein by flow cytometry. The summarized data are shown. Assays were performed in triplicate. The dashed horizontal lines indicate the value of BA.2. Statistically significant differences versus each parental S protein and those between BA.2 were determined by two-sided Student’s *t* tests.

**(G)** S-based fusion assay in Calu-3 cells. The recorded fusion activity (arbitrary units) is shown. The dashed green line indicates the result of BA.2. The red number in each panel indicates the fold difference between BA.2 and the derivative tested at 24Dh post coculture. Assays were performed in quadruplicate. Statistically significant differences versus BA.2 across timepoints were determined by multiple regression. The familywise error rates (FWERs) calculated using the Holm method are indicated in the figures.

**(H and I)** Neutralization assay using BA.2 breakthrough infection (BTI) sera **(H)** and BA.5 BTI sera **(I)**. Neutralization assays were performed with pseudoviruses harboring the S proteins of B.1.1, BA.2, EG.5.1 and BA.2.86. The following sera were used. I, J Convalescent sera from fully vaccinated individuals who had been infected with BA.2 after full vaccination (9 2-dose vaccinated and 4 3-dose vaccinated). 13 donors in total) (**H**), and BA.5 after full vaccination (1 2-dose vaccinated donors, 15 3-dose vaccinated donors and 1 4-dose vaccinated donor). 17 donors in total) (**I**). Assays for each serum sample were performed in triplicate to determine the 50% neutralization titer (NT_50_). Each dot represents one NT_50_ value, and the geometric mean and 95% confidential interval (CI) are shown. Statistically significant differences were determined by two-sided Wilcoxon signed-rank tests. The P values versus B.1.1 or EG.5.1 are indicated in the panels. The horizontal dashed line indicates the detection limit (40-fold). Information on the convalescent donors is summarized in **Table S2**.

