## Supplementary figures and images for "Virological characteristics of the SARS-CoV-2 BA.2.86 variant"

### Figure S1

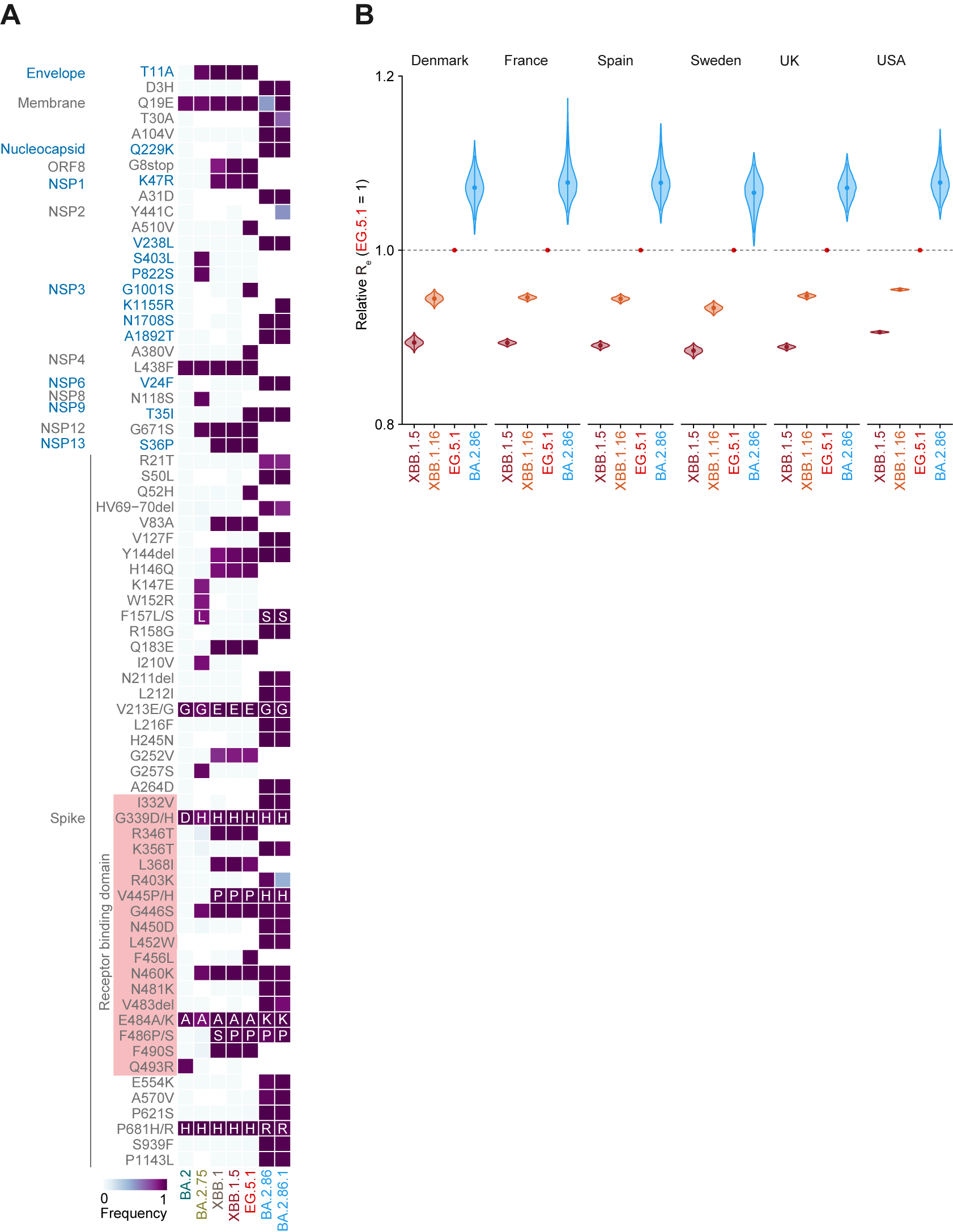

### Figure S2

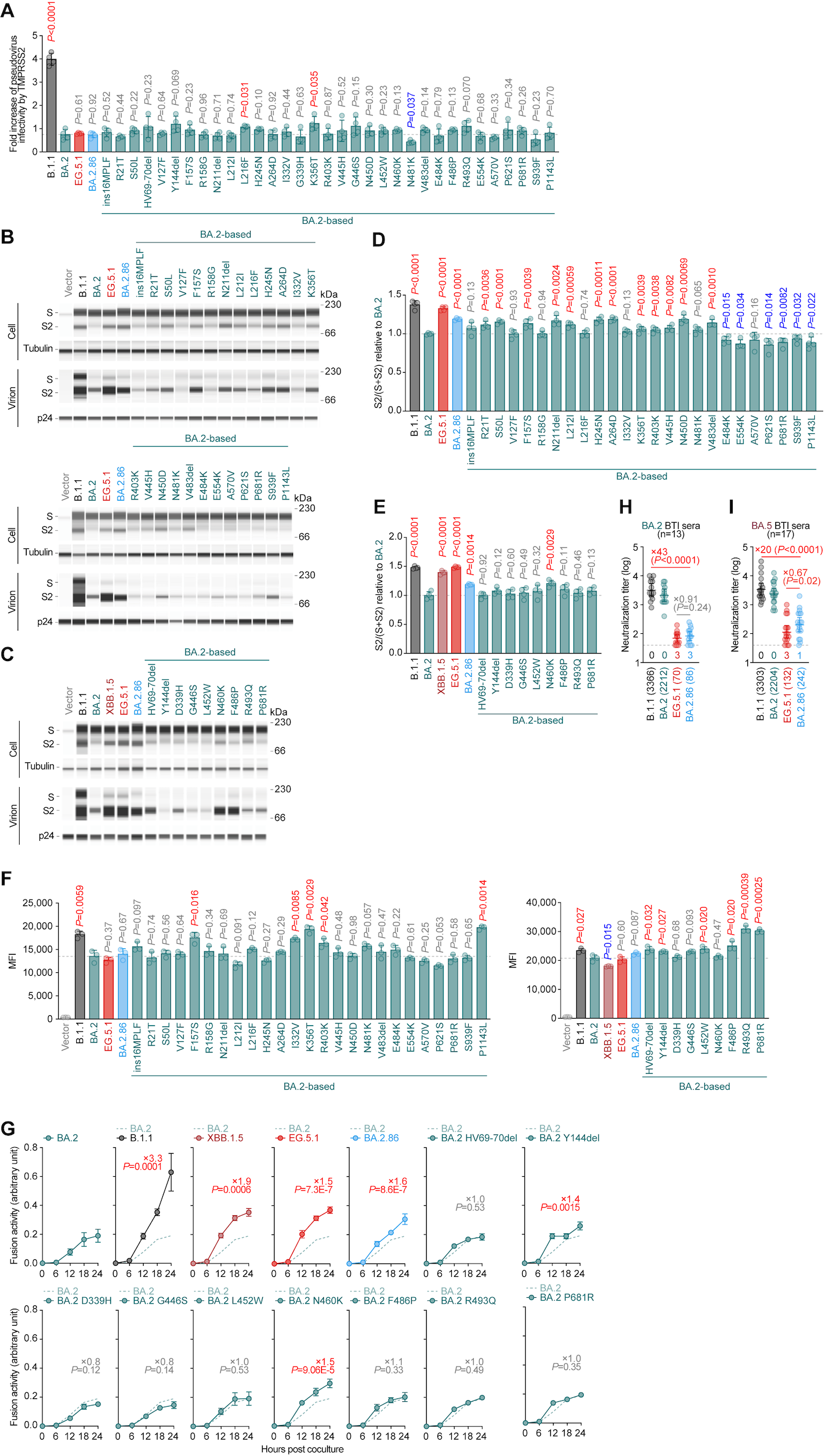
